# A Functional Data Analysis Approach to RuBisCO Engineering

**DOI:** 10.1101/2023.06.14.544690

**Authors:** Adam C. McCormack

## Abstract

Mitigating and adapting to climate change requires carbon emission control and effective technologies for drawing greenhouse gases from the atmosphere. Here we propose an effective strategy for guiding the rubisco engineering problem, which seeks to improve photosynthesis and carbon sequestration in crops by minimizing photorespiration, a major impediment to crop yields. Photorespiration occurs when rubisco oxygenates rather than carboxylates, thus reducing carbohydrate synthesis to eliminate toxic byproducts. Most plants, including most agricultural crops, exhibit a C3 photosynthetic pathway and have not developed adaptive mechanisms to combat rubisco’s tradeoff. The span of low activity rubiscos opens the possibility of engineering less productive crops to express higher activity enzymes. The main experimental challenges in bypassing rubisco’s biochemical limitations include its molecular size and complexity, lack of empirical data, and costs of acquiring new data, but recent advances in machine learning and data analytics could allow us to overcome these challenges. In particular, we propose a novel computational approach to inform experimental research into rubisco engineering by employing recently developed techniques from functional data analysis. We show that the separation between high and low activity enzymes can be modeled within the sequence space of rubisco’s primary structure, and we further discuss how our approach can guide deeper investigation into the rubisco engineering problem. Future empirical success would simultaneously address major global issues such as rising atmospheric CO_2_ levels and food insecurity.

## 1 Introduction

The evidence for anthropogenically driven climate change is now overwhelming. We observe its impact on many aspects of the human condition, including food security, drought, wildfires, and flooding [1–3]. Proposals to regulate the emission of causative agents, such as carbon dioxide, nitrous oxide, methane, and other trace gases via policy enforcement are abundant yet ineffective considering that the average global concentration of these agents continues to increase while its impact on human life and the environment becomes more severe each year [4, 5]. Fossil fuel CO_2_ emissions in particular present a unique challenge such that an estimated 20-25% of the released volume could remain in the atmosphere up to a millennium, further producing positive feedback in the destabilization of other natural carbon sinks [3]. This information implies that an effective strategy for mitigating climate change must include the development and application of technologies for drawing CO_2_ out of the atmosphere.

Numerous ideas have been proposed to facilitate the development of impactful carbon capture technologies, and some approaches have even been discussed for decades. Agrigenomics is an important category of proposed strategies with potential for high impact, however, it has received little attention until recently [6]. One attractive aspect of a plant-based genomic strategy is that, alongside carbon capture, it simultaneously offers the possibility of progress on another pressing issue: ensuring food security by increasing agricultural crop yields [7, 8]. Recent empirical advances in agricultural gene editing technologies have shown promising results [9–11], but we have yet to observe viable results specifically for plant rubisco engineering. Researchers have been successful in heterologous expression of cyanobacterial components within tobacco, but in all cases the resulting organism was nonviable at atmospheric CO_2_ levels [12–14]. More prudent approaches must therefore be established to increase crop yield at ambient atmosphere via rubisco engineering.

Three primary strategies have been proposed for plant rubisco-focused synthetic biology approaches to carbon capture: 1) designing new-to-nature rubisco enzymes with superior activity, 2) transplanting rubisco components from highly active organisms into plants with lower activity, and 3) bypassing rubisco entirely via introducing novel metabolic pathways as an alternative to the Calvin-Benson-Bassham cycle [15]. Rubisco’s molecular complexity and associated costs of enzyme-related experimentation pose significant challenges to the former two strategies, but recent advancements in high-throughput computation could allow us to overcome these obstacles. More specifically, computational techniques such as machine learning have the potential to direct empirical research into rubisco engineering such that resource costs are minimized, and the likelihood of experimental success given rubisco’s complexity is maximized. Machine learning applications in the field of enzyme engineering are at the forefront of scientific advancement, and continuous development and success in this space have been demonstrated in the literature [16–18]. This establishes a central scientific motive to develop novel computational techniques capable of informing experimental research into rubisco engineering.

The rubisco enzyme is naturally present in four distinct isoforms, forms I, II, III, IV, with form I being the most abundant and being expressed in both plants and bacteria. Form I is also the only isoform composed of two distinct structural subunits, the catalytic large subunit (LSU) and non-catalytic small subunit (SSU). It has been established that SSU structure affects kinetic activity within the form I group [19], and therefore, to best inform plant rubisco engineering with computation, we seek to build a machine learning framework that incorporates the sequences of both subunits and predicts enzyme function, e.g., kinetic and binding activities. However, construction of a training set is limited by the scarcity of molecules for which both the complete sequences and kinetic activity are known. At present, a useful sequence-kinetic training set can only be constructed using LSU sequences, which may suffice at the current stage considering that the LSU contains the catalytic site. Furthermore, a decision algorithm must be identified that works well with the limited size of the training data and the abundance of features needed to encode protein sequences.

Positive results have recently been reported for Embryophyte LSU sequence-kinetic regression [20], and phylogenetic interrogation of bacterial and archaeal rubiscos has also proven successful within the Form II-III phenotypic stratum [21, 22]. Here we introduce the Multilevel Orthogonal Subspace (MOS) feature engineering algorithm, which constructs separable feature vectors for functionally related class data. The separability of MOS features facilitates classification tasks on topologically complex domains and has been shown to produce mathematically robust and explainable class predictions without knowledge or assumptions on the underlying residual distribution [23, 24]. Fundamental results on the statistical and biological importance of functional spaces for protein engineering have proven insightful for simpler molecules [25, 26], but the method we employ is intrinsically novel. Our approach also contrasts with more popular computational tools being applied in this field, such as neural networks and gaussian process models. We demonstrate the efficacy of the MOS algorithm as applied to multiple rubisco sequence-function prediction protocols, and we benchmark it against previously published results where applicable. We finally discuss how our application can be used to direct the rubisco engineering problem, potential and necessary routes for future empirical studies, and any limitations of our ideas and strategies for instructing the proposed experimentation.

## 2 Materials and Methods

### 2.1 Computational Experiment Design

In this work, natural realizations of rubisco’s amino acid sequence and their function were modeled as a stochastic process to make biological predictions on the function of empirically unknown rubiscos. The novel MOS feature engineering algorithm was analyzed on its ability to construct separable features for rubisco sequence-function data, which was cross-validated using a binary support vector machine (SVM) decision algorithm. A sigmoid (SIG) transformation function was then estimated via Platt Scaling [27] to assign class posterior estimates to SVM output. The MOS+SVM+SIG framework was assessed on its reliability across two varying protein sequence-function prediction experiments. For the first experiment, the proposed framework was trained on empirically labeled LSU sequences and used to predict the kinetic activity of unlabeled LSU sequences. This procedure was cross-validated using C3 plant data as a control group and Embryophyte data, curated by Iqbal et al. [20], as a validation/comparison set. The ability of our algorithm to identify kinetically desirable rubisco enzymes was also assessed across higher order phenotypic strata. In the second experiment, SSU sequences were incorporated to predict functional and non-functional variants of SSU and LSU combinations, which was cross-validated using empirical data on *Synechoccus elongatus* PCC6301. The purpose of this work is to demonstrate our ability to guide the rubisco engineering challenge using novel computational techniques.

### 2.2 Curating Sequence-Function Training Sets

Amino acid sequences used in this study were obtained either as supplementary material from relevant literature [21] or from the UniprotKB online database [28] by querying ‘family:”RuBisCO large chain family”’ and ‘family:”RuBisCO small chain family”’ for the LSU and SSU, respectively. Enzyme kinetic activity data were collected from the literature [21, 29–31] and further curated with their respective taxonomic identifiers. Rubisco sequences collected from online resources were then matched to their corresponding kinetics using the species’ unique taxonomic key. Protein sequences with unknown amino acid residues were allowed in our study, and multiple sequences belonging to the same species were also included by multiply assigning kinetic labels. All amino acid sequences were numerically encoded using the Clustal Omega algorithm (ver. 1.2.4) under default parameters to construct a single Multiple Sequence Alignment (MSA) for all sequences in the training and validation datasets [32]. The MSA was cleaned at varying gap resolutions (GR), where MSA sequence indices with a columnar gap percentage greater than the GR percentage are removed from all sequence vectors. Individual sequences were ultimately vectorized using the five amino acid factors identified by Atchley et al. [30], effectively quintupling the length of each MSA sequence. A neutral factor score of zero was introduced for any remaining gapped positions or unknown amino acid residues present in the MSA sequences.

### 2.3 MOS Feature Construction

The MOS algorithm constructs *ℓ* levels, or nested subspaces, orthogonal to a *k*-eigenvalued approximation of the protein sequence vector covariance spectrum, which form a one-to-one mapping function 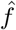 that transforms amino acid sequence vectors into their respective MOS feature vectors 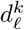. The MOS algorithm is trained using the majority class data of a class-imbalanced dataset, and the mapping function 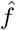 may be considered a filter that transforms amino acid sequences divergent from the majority class covariance spectrum into a new feature space with likelihood guarantees on separability. The magnitude of MOS features 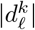 is bounded by the decay rate of the covariance spectrum eigenvalues *λ*_*j*_, and the existence of a natural probability measure on MOS feature posterior separability has been mathematically proven by Kon and Castrillón-Candás [23, 24]. Construction of MOS features therefore allows for robust hypothesis testing of functional class data on topologically complex domains, which may be achieved by choosing a significance level *α* and constructing an estimate for the following probability measure:

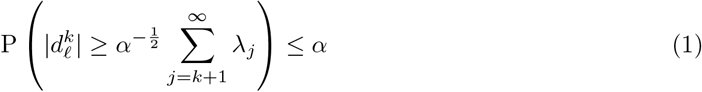

Equation 1 states that the magnitude of MOS features is bounded by the residual space of the covariance eigenspectrum, i.e., the leftover portion of an infinite Karhunen-Loève representation that results after approximating, or truncating, to *k* terms [23, 24]. We use the SVM posterior function to estimate this probability measure by finding the residual separation between two classes. As we demonstrate later, choosing the significance level *α* = 0.05 allows us to classify majority class observations with 95% confidence.

### 2.4 Statistical Analyses

For the first experiment, kinetic parameter labels for homologous sequence vectors were averaged after MSA encoding, thus guaranteeing the uniqueness of each feature vector. Sequence vectors were labeled as class 0 or class 1 according to the specific hypothesis being tested, with class 0 being the majority class in all experiments. The minority class was oversampled in the first experiment using a variation of the Synthetic Minority Oversampling Technique (SMOTE) [33] in which artificial data were generated by kriging synthetic sequence vectors from the covariance spectrum of natural sequence vectors [24]. The MOS+SVM+SIG framework was then cross-validated on artificially balanced class data using a leave-one-out approach. For the second experiment, random undersampling of the majority class was used with repeated cross-validation due to the drastic imbalance of classes. Posterior probability estimates of class labels P(*x*_*i*_ ∈ Class 1 | *k, ℓ, X*), henceforth 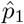, were used to model the equation 1 probability measure with significance level *α*. Probabilistic model calibration is quantified using the Brier Score (BS) metric:

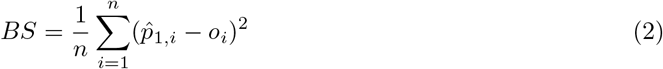

The BS calibration metric describes the accuracy of probabilistic predictions on a labeled training set of size *n*, where the continuous variable 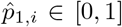 [0, 1] is the minority class posterior probability and the discrete variable *o*_*i*_ ∈ {0, 1} is the true class label of the *i*^*th*^ observation. The continuous metric value *BS* ∈ [0, 1] denotes perfect accuracy at *BS* = 0 and perfect inaccuracy at *BS* = 1 [34].

## 3 Results

### 3.1 Determining a Kinetic Threshold for Highly Active Rubiscos

To perform classification on enzyme activity, continuous kinetic parameter labels were partitioned into discrete classes. Binary labels *high* (class 1) and *low* (class 0) were used to describe the desirability of an enzyme’s kinetic activity, and a discretization threshold was selected to assign class labels for hypothesis testing. As we discuss later, clearly defining the hypothesis is critical for the appropriate generation and interpretation of results by our proposed framework. We formally state the principal hypothesis for the first experiment as follows:

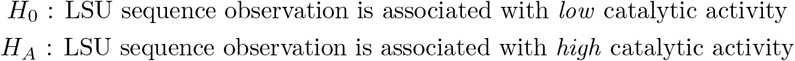

A kinetic discretization threshold was determined by comparing agricultural C3 crop rubisco kinetics with those of non-agricultural C3 plants that express best-known enzyme activity. A sample dataset shown in table 1 highlights the kinetic differences between these C3 subgroups. From the data it appears that the CO_2_ michaelis-menten constant *K*_*C*_ and molecular turnover rate *k*_*cat,C*_ of the best-known C3 species are roughly double those of agricultural C3 species, with a slight increase in CO_2_ substrate specificity *S*_*C/O*_. This fact suggests that it may be possible to improve the rate of carboxylation in agricultural C3 crops by transplanting rubisco genes from the best-known C3 species, and therefore it would be advantageous to develop a method for identifying other *high* activity enzymes out of the empirically unknown pool of plant species.

**Table 1:** Sample rubisco kinetic observations demonstrating the difference in enzyme kinetic parameters between agricultural and best-known C3 plants. From the data it appears there exists significant kinetic differences between C3 subgroups, and therefore it may prove insightful to study plant rubisco transgenics using donors closely related in phenotype.

| Agricultural C3 Plants |  |  |  | High Activity C3 Plants |  |  |  |
| --- | --- | --- | --- | --- | --- | --- | --- |
| Species | $K_C(\mu\text{M})$ | $k_{cat,C}(\text{s}^{-1})$ | $S_{C/O}$ | Species | $K_C(\mu\text{M})$ | $k_{cat,C}(\text{s}^{-1})$ | $S_{C/O}$ |
| *N. tabacum* | 10.7 | 3.4 | 82 | *A. latifolia* | 21.0 | 5.8 | 105 |
| *S. oleracea* | 11.0 | 2.7 | 92 | *P. maritima* | 20.8 | 5.4 | 106 |
| *G. max* | 13.2 | 2.5 | 100 | *P. distans* | 22.2 | 5.4 | 104 |
| *I. batatas* | 12.0 | 2.5 | 98 | *A. stolonifera* | 25.3 | 5.2 | 105 |
| *A. sativa* | 10.8 | 2.3 | 100 | *P. limmonii* | 28.1 | 5.2 | 102 |
| *T. aestivum* | 11.3 | 2.2 | 100 | *L. giganteum* | 31.2 | 5.1 | 108 |
| *C. maxima* | 9.0 | 2.2 | 98 | *F. pratensis* | 23.1 | 5.1 | 106 |
| *C. arabica* | 11.0 | 2.1 | 99 | *D. danthoniodes* | 22.3 | 4.5 | 108 |
| *O. sativa* | 8.0 | 2.1 | 93 | *C. murale* | 23.8 | 4.4 | 109 |

Discretization thresholds were chosen around the example in table 1 since this data addresses one of the main rubisco engineering strategies: transplanting more active rubisco components into crop species of lesser activity. Two different thresholds were chosen to create partitions with varying class distribution, which we refer to as control groups A and B. Discretization according to the selected thresholds result in a class label of *high* for C3 plants only, which is important for our application since C3 plant expression of non-C3 rubiscos from higher order phenotypes has been shown to be nonviable at ambient atmosphere [12–14]. As we initially demonstrate, the discretization threshold affects framework performance but also ensures that *high* class predictions belong to the appropriate target phenotype. Depicted in figure 1 is a visual representation of how the discretization threshold affects class imbalance and MOS training set size within the C3 subgroup.

**Figure 1:**
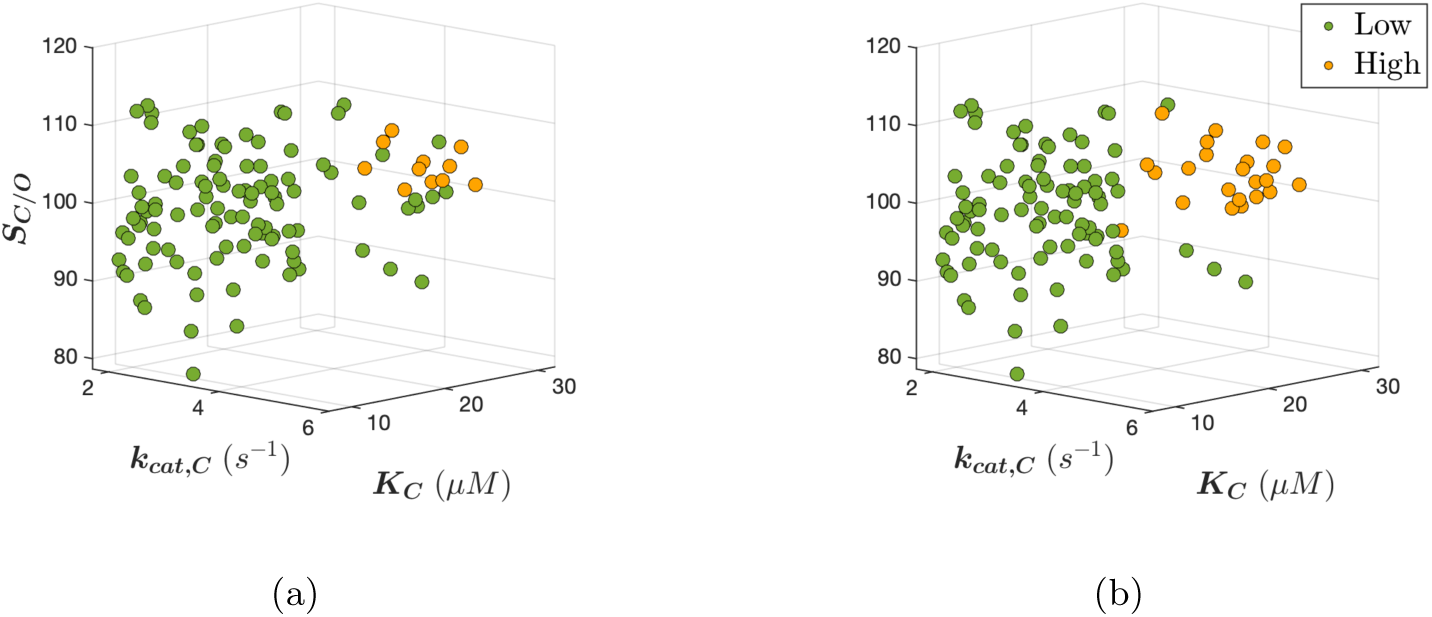
A depiction of how the kinetic discretization threshold affects training set size for the control groups. A total of *n* = 207 unique sequences were identified for the C3 plant phenotypic stratum using the natural encoding scheme at 50% GR. (a) The control group A threshold *k*_*cat,C*_ *>* 4*s*^*−*1^; *K*_*C*_ > 20*μM*; *S*_*C/O*_ *>* 100 resulted in class sizes *n*_1_ = 20 and *n*_0_ = 187. (b) The control group B threshold *k*_*cat,C*_ *>* 3.5*s*^*−*1^; *K*_*C*_ *>* 18*μM* ; *S*_*C/O*_ *>* 95 resulted in class sizes *n*_1_ = 39 and *n*_0_ = 168.

For the partitions depicted in figure 1, green *low* activity sequences were used to train the MOS transform function, which was then applied to all sequence vectors to obtain their corresponding MOS feature vectors. It may appear from figure 1 that the classes are clustered and wellseparated within the kinetic fitness space, however, the challenge lies in finding separation within the amino acid sequence space. The MOS algorithm attempts to find this separation within the residual spaces of the amino acid sequence covariance spectrum to engineer features with likelihood guarantees on separability, thus allowing for easier classification using a standard decision algorithm such as SVM. F-score classifications performance is demonstrated in figure 2 for posterior models constructed from a binary SVM under both control group thresholds. Performance was cross-validated on balanced class data after kriging synthetic data from the minority class covariance spectrum.

**Figure 2:**
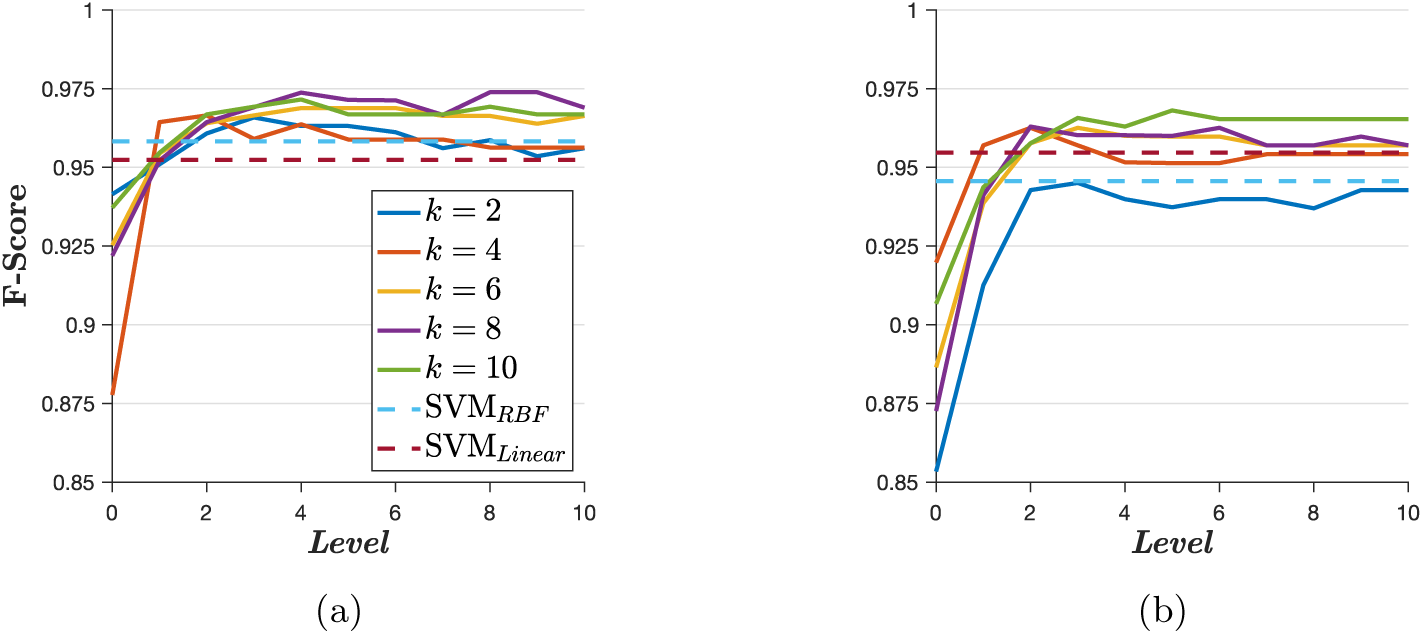
Cross-validated F-score metrics for posterior models constructed from control groups A and B at 50% GR. (a) Control group A posterior classification performance across the MOS hyperparameter space. The multilevel filter appears to converge at higher levels. (b) Control group B posterior classification performance. The F-scores appear more sparse at higher residual levels and with slightly less accuracy than control group A.

It was observed that classification performance for control group A was slightly higher than control group B, with some improvements from SVM without MOS features. When supplied with functional data, this framework effectively operates as an anomaly detection filter. The standard F-score accuracy metric was initially reported since class data was artifically balanced using SMOTE, however, it does not inform us on the quality of calibration achieved by the probabilistic model [35]. Further analysis of the control group posterior probability distributions using a proper scoring rule, such as the BS calibration metric, is necessary to reach the appropriate conclusions on posterior model reliability.

### 3.2 Calibrating the Posterior for Enzyme Activity Predictions

This section covers an analysis of the posterior distributions to determine which discretization threshold, hyperparameters, and phenotypic training stratum result in the best performance. Model calibration was quantified by calculating the BS from the predicted posteriors, and the best hyper-parameters were selected by minimizing the BS. Shown in figures 3a and 3b are the control group BS metrics across the MOS hyperparameter space, where it becomes clear that the control group A partition results in better overall calibration. However, further stratifying the BS according to class label 0 or 1, as shown in figures 3c and 3d, reveals that the control group A partition also results in better calibration for both classes. The minimum BS was achieved using hyperparameters {*k* = 6, *ℓ* = 4}, however, the model obtained using {*k* = 8, *ℓ* = 4} resulted in a 0.001 difference from the minimum BS but with a significant increase in F-score, as reported in figure 2a. The latter set of hyperparameters was therefore determined to produced the best calibration.

**Figure 3:**
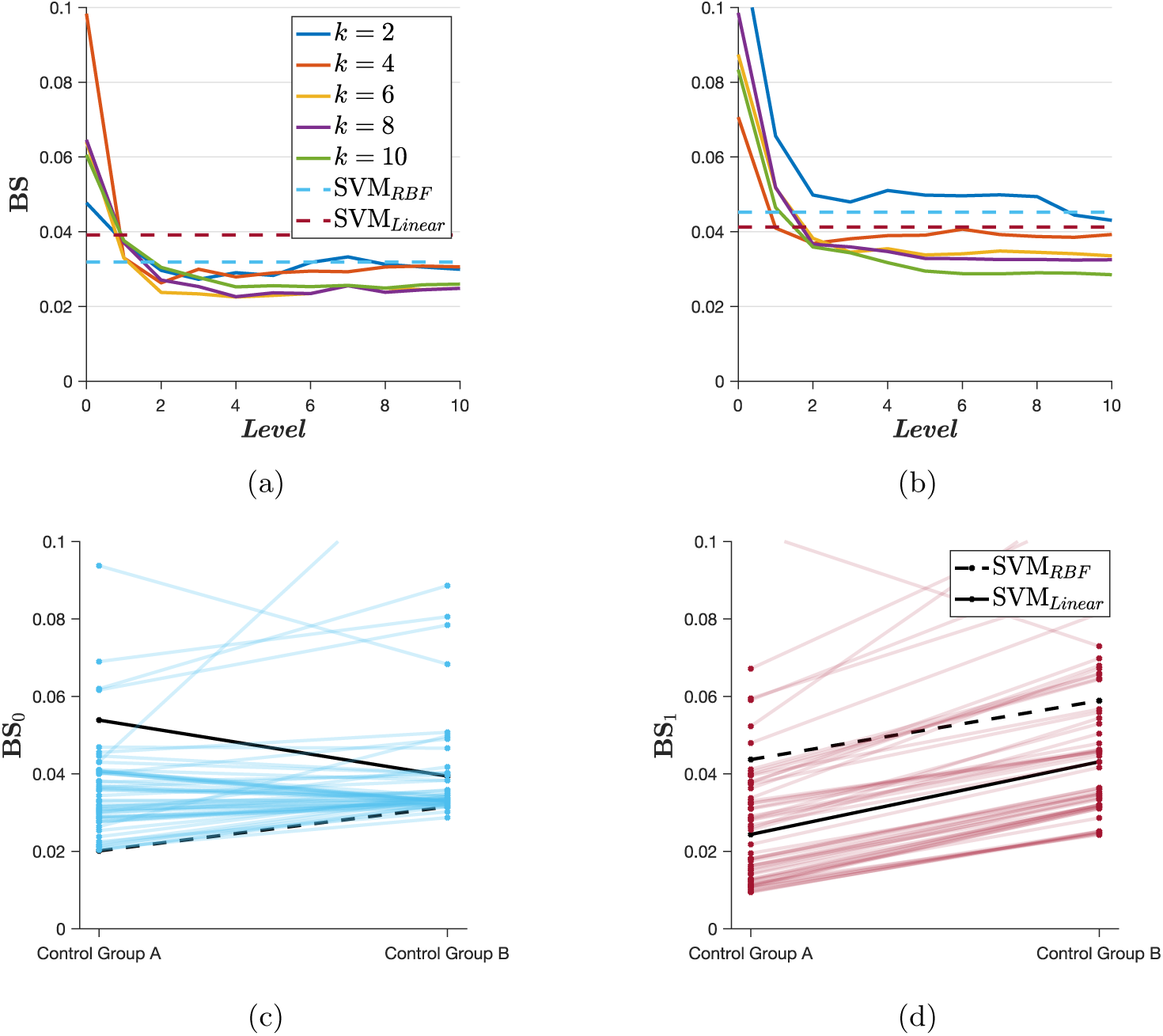
Cross-validated BS metrics for control groups A and B at 50% GR. (a) BS progression for control group A across the MOS hyperparameter space. (b) BS progression for control group B. It is clear that control group B partition achieves less overall calibration. (c) Slopegraph for stratified BS of class 0 across both control groups and all MOS hyperparameters. For the majority class, it was observed that SVM*_RBF_* without MOS features converged to the same calibration as MOS+SVM. (b) For the minority class, it is clear that MOS features produce a better calibration when compared with SVM*_RBF_* and SVM*_Linear_*.

Visual inspection of the predicted posterior distribution conveys the translation from mathematical theory to empirical data analysis. Recall that the equation 1 probability measure describes the theoretical posterior separability of MOS features, which is estimated by the SVM posterior function. We demonstrate this concept in figures 4a and 4b using models with the worst and best calibration, respectively, according to previous analyses. It is important to first calibrate model hyperparameters, as the *α*-threshold may be used as the decision boundary for classifying *high* or *low* activity sequences. For well-calibrated models and applications in which the associated costs of a false positive prediction are relatively low, it becomes beneficial to use the *α*-threshold for assigning class labels to empirically unknown data. This is because statistical significance can be assigned within our hypothesis testing framework while simultaneously maximizing the True Positive Rate (TPR).

**Figure 4:**
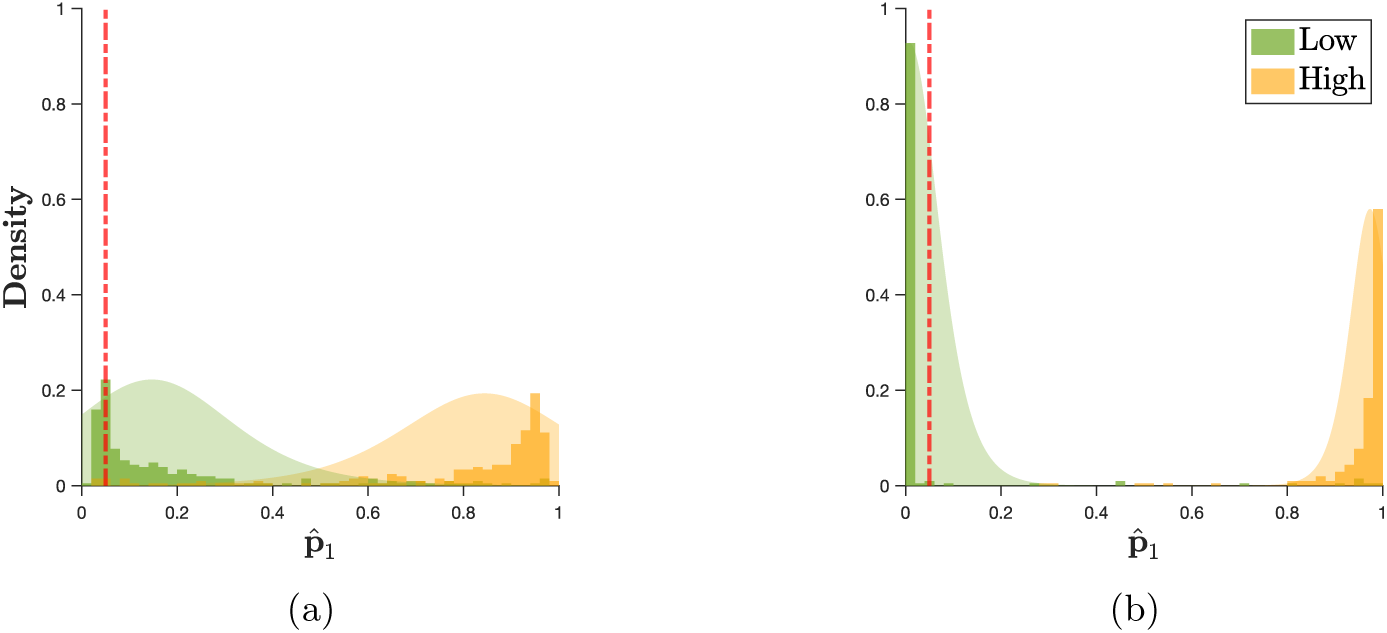
Cross-validated posterior distributions for control group A, with the significance level *α* = 0.05 indicated by the dashed red line. (a) Posterior distribution for the worst-calibrated model at *k* = 4 eigenvalues and *ℓ* = 0 levels. There appears to be good separation along the default classification threshold at 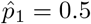, however, a large portion of class 0 predictions remain above the *α*-threshold. (b) Posterior distribution for the best model at *k* = 8 eigenvalues and *ℓ* = 4 levels. We should expect that well-calibrated MOS filters map 95% of the class 0 posterior estimates fall below the *α* = 0.05, as described by equation 1.

Before making predictions on unlabeled LSU data, a final analysis of the purely supervised framework was conducted in which we deviate from the control group by amplifying evolutionary variance within the training set data. This was accomplished by increasing the GR and including higher order phenotypic data in the observation set. Recall that the kinetic discretization thresholds are unique to C3 plants, meaning that including higher order phenotypes in the observation set does not affect the variance of class 1 data but does contribute to the variance of class 0 data, which is used to train the MOS mapping function. Based on the results in table 2, it is evident that higher order phenotypic variance increases the calibration of the MOS covariance filter for detecting anomalous *high* activity rubisco sequences.

**Table 2:** Cross-validated BS calibration metrics using the control group A discretization threshold at increased GR and higher order phenotypic strata. The purpose of these trials is to examine the effects of evolutionary variance on model performance, where the C3 plant stratum at 50% GR represents the least training variance and the Forms I, II, III stratum at 99% GR represents the greatest training variance.

| Training Strata | Gap Resolution |  |  |  |
| --- | --- | --- | --- | --- |
|  | 50% | 75% | 95% | 99% |
| **C3 plants** | 0.0225 | 0.0217 | 0.0223 | 0.0218 |
| **Embryophyta** | 0.0175 | 0.0181 | 0.0182 | 0.0180 |
| **Eukaryota** | 0.0148 | 0.0165 | 0.0170 | 0.0163 |
| **Form I-B** | 0.0151 | 0.0186 | 0.0186 | 0.0181 |
| **Form I** | 0.0119 | 0.0149 | 0.0159 | 0.0158 |
| **Forms I, II, III** | 0.0106 | 0.0126 | 0.0131 | 0.0131 |

### 3.2 Incorporating Unlabeled LSU Sequence Data

In the previous section, we cross-validated the ability of our posterior framework to assign class posterior estimates to labeled data in a purely supervised approach. In this section, we analyze the posterior function’s ability to generalize to unlabeled Embryophyte data curated by Iqbal et al [20], and we benchmark our prediction set against theirs. First, the control group A discretization threshold was applied to Iqbal et al.’s regression results to obtain the relevant discrete class labels. The MOS+SVM+SIG framework was then applied to the raw sequences to obtain posterior class predictions using the Forms I, II, III phenotypic stratum at 50% GR, since this data was previously reported to produce the best calibration. The validation set posterior distribution and associated training set classification metrics are reported in figure 5. For the selected training stratum, a minimum BS was achieved at both {*k* = 6, *ℓ* = 2} and {*k* = 8, *ℓ* = 2}. The latter set of hyperparameters maximizes the cross-validated TPR and was therefore selected to assign posterior estimates to the validation set.

**Figure 5:**
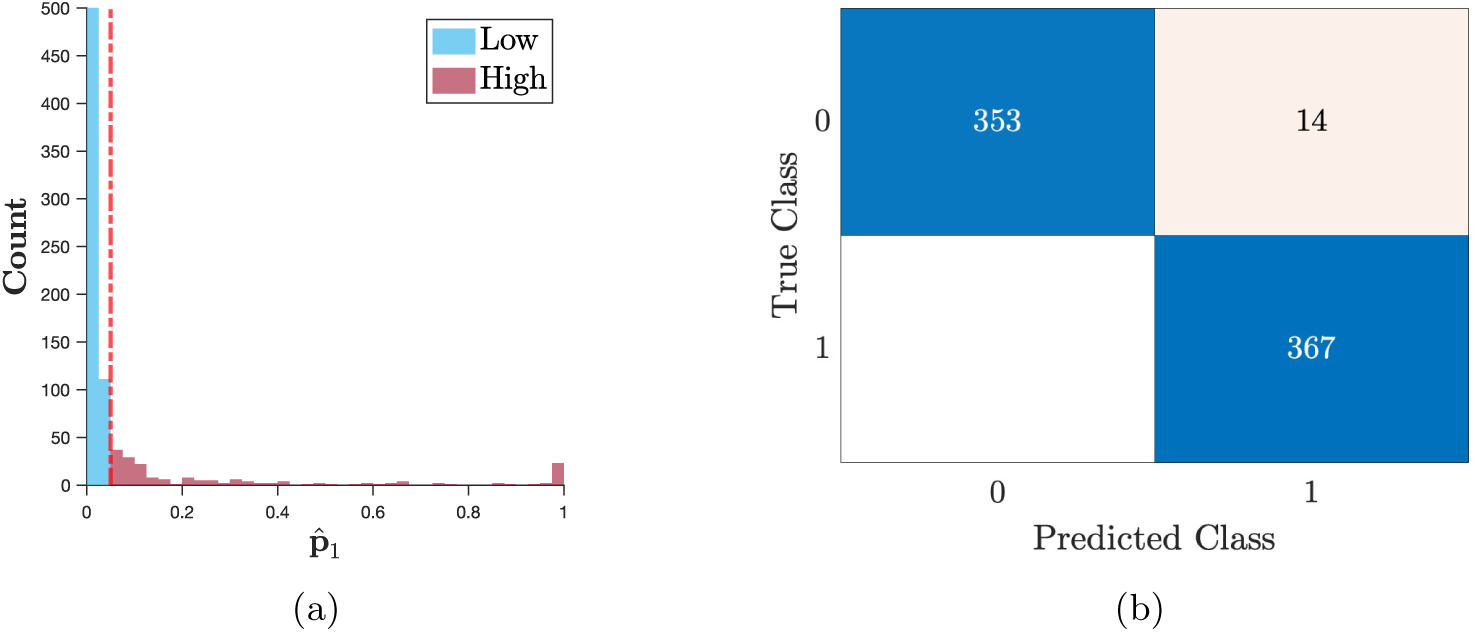
Posterior estimation results for the unlabeled Embryophyte validation set of size *n*_*u*_ = 14, 456 and associated classification metrics for the labeled training set. (a) Predicted posterior count distribution resulting in 98.7% of the validation set with 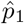 below the *α*-threshold and predicted class distributions 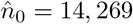, 269 and 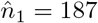. (b) Cross-validated *α*-threshold classification metrics for the Forms I, II, III training stratum at 50% GR, with empirical class distribution *n*_0_ = 367 and *n*_1_ = 20 according to the control group A threshold. Training set class distributions were artificially balanced using SMOTE to obtain a training set of size *n* = 734.

According to the calibrated posterior function and *α*-threshold decision boundary, our framework identified 187 unique *high* activity sequences, of which 184 belong to C3 plants and 3 belong to C4 plants. These results make sense considering the natural phenotypic distribution of *high* activity sequences within the empirical observation data. The control group A threshold was applied to the gaussian process results reported by Iqbal et al. to identify 135 unique *high* activity rubisco sequences from their prediction set. Interestingly, there is predictive agreement for only 22 species out of both prediction sets, which indicates a fundamental difference in the underlying behavior of both predictive techniques. At present, it may be prudent to devote experimental resources to predictions upon which both models agree.

### 3.4 Incorporating SSU Sequence Data

We conclude our methodology with a brief section on a different rubisco engineering application. SSU sequences were excluded from the first experiment due to the lack of available sequences for the *high* activity class, but, by slightly rephrasing our hypothesis, we can use the MOS+SVM+SIG framework to investigate another important question involving SSU data. A singular rubisco enzyme is a macromolecular complex composed of multiple LSU and SSU protein subunits, and its construction requires a series of intermediate protein-protein interactions [36]. To engineer such a system, it would be useful to develop a statistical model for predicting which combinations of LSU and SSU sequence variants may produce an active enzyme. This second experiment differs from the first such that the magnitude of kinetic activity is no longer of immediate interest. To direct the design of synthetic enzymes using novel combinations of natural subunit sequences, a posterior estimate was used to test the following hypothesis:

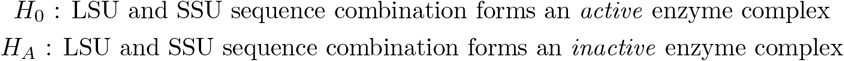

It is an important point that under the alternative hypothesis we make no claims about the physical binding of protein subunits, yet this fact is implied under the null hypothesis. A subunit combination belonging to the *inactive* class may or may not physically bind, but in both cases the alternative hypothesis states that the combination expresses no kinetic activity. This notion indicates the importance of training set construction, which impacts the appropriate conclusions drawn from our framework. The second experiment was initiated by bootstrapping posterior estimates on cyanobacterial form I subgroup B rubisco data belonging to *S. elongatus* PCC6031, for which 64 observations of *inactive* sequence combinations were identified from relevant literature [30, 31]. The majority class training set consisted of 15,841 *active* combinations of LSU and SSU sequences across all Form I taxonomic lineages. Results for this experiment are displayed in figure 6 below.

**Figure 6:**
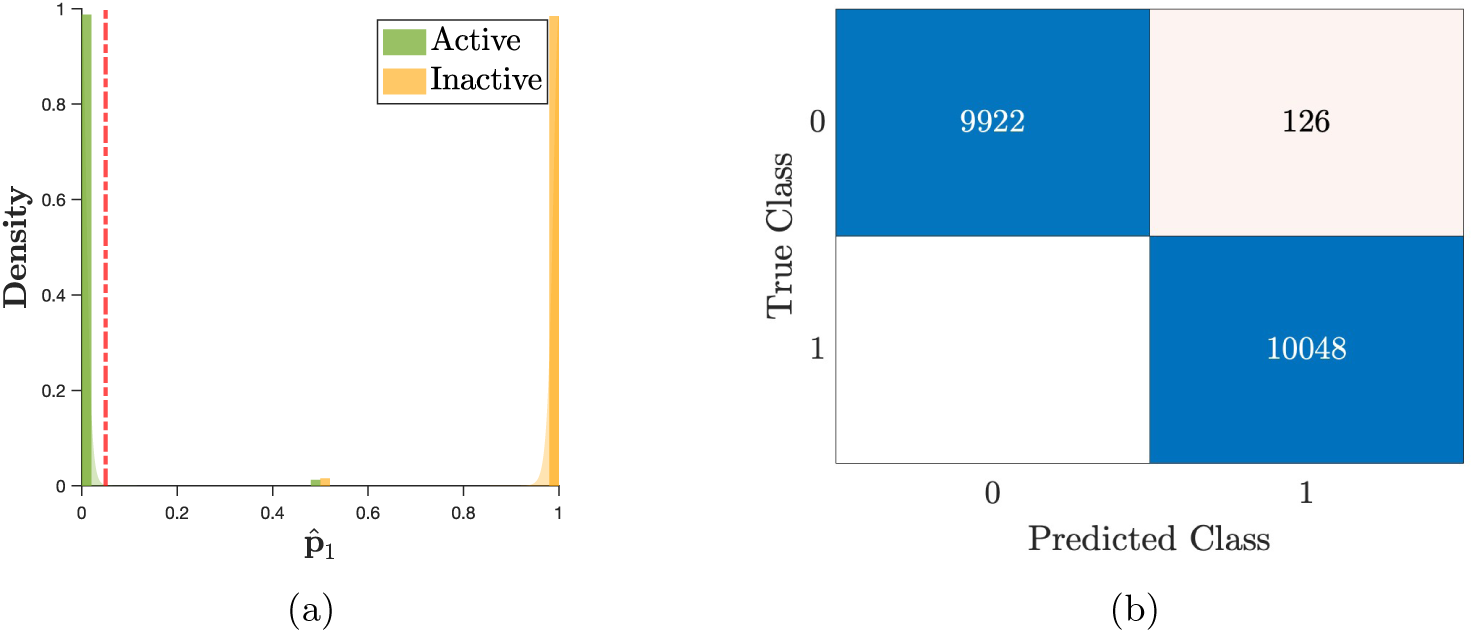
Posterior distribution and *α*-threshold prediction results for the second experiment using the natural encoding scheme at 50% GR with hyperparameters *k* = 2, *ℓ* = 4. (a) Class posterior distribution of the bootstrapped posteriors obtained after random undersampling of the majority class. Greater calibration for this experiment is obvious compared to the distribution from figure 4b. (b) Cross-validated *α*-threshold prediction results across all iterations of cross-validation.

The bootstrapping procedure for this experiment was conducted as follows: 157 iterations of cross-validation were performed after randomly undersampling the majority class, resulting in a class 0 sample of size 64 and holdout set of size 15,777. The MOS algorithm was trained on the holdout set and used to generate features for the balanced class data, which were then used to crossvalidate an SVM+SIG posterior model. The leave-out test observation from cross-validation was used as a singular bootstrap sample, thus creating 64 bootstrap estimates per iteration and ensuring a minimum of 10,000 bootstrap samples. From this procedure we observed that the MOS+SVM+SIG framework achieved significantly higher calibration than the previous experiment, which is likely due to the greater MOS training set size. This experiment demonstrates the robustness of our framework in identifying functional combinations and anomalous single-point mutations in the rubisco subunit sequences.

## 4 Discussion

The proposed posterior modeling framework presents an effective strategy for guiding the rubisco engineering challenge, which seeks to increase carbon sequestration and biomass yields in agricultural crops by reliably identifying specific enzymes that exhibit superior rubisco kinetics. Our method was shown to generate robust functional predictions for various protein sequence manipulations that would be useful for guiding experimental enzyme engineering research in the form of data-driven hypothesis tests and class posterior estimates. A useful next step for experimental research would be to empirically validate the posterior predictions from our first experiment, though it would also be worthwhile to obtain missing SSU sequences for empirically measured enzymes and recreate the first experiment with SSU data.

A drawback to our approach is that we have not yet identified a statistically robust method for designing completely synthetic enzymes, however, experimentalists interested in designing new-to-nature rubiscos may generate novel combinations of natural LSU and SSU sequences and utilize the approach from our second experiment to determine enzyme viability. The SMOTE method from our first experiment utilizes a naive approach to synthetic sequence generation by inducing a continuous normal distribution across the discrete amino acid sequence space. While this SMOTE procedure is useful and valid for model calibration, the synthetic sequences generated here are most likely non-viable in nature. Chowdhary and Najm have identified an iterative Bayesian approach to sampling the functional covariance spectrum [37], which may be useful for generating completely synthetic enzymes. This generative technique could establish the superiority of MOS+SVM+SIG over SVM+SIG (without MOS features), since the latter algorithm is non-generative.

Another drawback to our approach is the lack of understanding of agricultural rubisco engineering. It is likely that the rubisco engineering challenge becomes a metabolic systems engineering challenge once genes of highly active enzymes are successfully transplanted into less active species. This notion is further admitted by previous transgenic studies that were carried out across large evolutionary distances [12–14]. We can propose that experimentation be conducted in which rubisco transgenic modification is performed across shorter evolutionary distances, i.e., within the C3 plant phenotypic subgroup. For example, according to the data in table 1, *P. distans* exhibits double *K*_*C*_ and *k*_*cat,C*_ with slight improvement in *S*_*C/O*_ when compared to *S. oleracea*. Transgenically modifying *S. oleracea* with the rubisco genes of *P. distans* would allow us to confirm our understanding of current metabolic models of photosynthesis and answer the question as to whether or not doubling rubisco’s carboxylation parameters results in the same increase in overall rate of photosynthesis and plant biomass yield. This approach would also allow us to identify downstream bottlenecks in the greater metabolic pathway.

It is a surprise that such experimentation has not yet been conducted, however, the difficulty behind this proposed experimentation arises from the fact that the SSU is coded in the nuclear genome and the LSU is coded in the chloroplast genome [38]. The associated transgenic experimentation may incur high experimental costs, but it would also offer valuable insights into the effects of rubisco transgenics on photosynthetic metabolism. Transgenic plant germlines would also need to be carefully engineered to ensure that transplanted genes proliferate throughout all plant cells. Based on the results presented in this paper, the proposed MOS+SVM+SIG framework may be used to mitigate the loss of resources that could arise from investigating undesirable species and enzymes. It is also possible that the proposed framework would be useful for analyzing the entire C3 photosynthetic metabolic pathway once appropriate data on C3 metabolic parameters have been obtained after successful LSU and SSU transgenic modification.

## Acknowledgments

I would like to thank colleagues Dr. Mark Kon and Dr. Julio Castrillόn-Candás for their useful discussion on the mathematics and implementation of the MOS algorithm. I would also like to thank Dr. Charles DeLisi for giving me the idea to use machine learning as a strategy for guiding the rubisco engineering challenge.

## Author Contributions

AM performed the methodology development, data processing and curation, computational experimentation, and relevant statistical analyses for this project. AM drafted all versions of this manuscript in its entirety.

## Data Availability

All associated results and data, both raw and processed, will be made available upon publication of this manuscript in a peer-reviewed journal. Source code for implementing the MOS algorithm is publicly available at https://github.com/jcandas/Multilevel-Change-Detection.

